# Complex population structure of the Atlantic puffin revealed by whole genome analyses

**DOI:** 10.1101/2020.11.05.351874

**Authors:** Oliver Kersten, Bastiaan Star, Deborah M. Leigh, Tycho Anker-Nilssen, Hallvard Strøm, Jóhannis Danielsen, Sébastien Descamps, Kjell E. Erikstad, Michelle G. Fitzsimmons, Jérôme Fort, Erpur S. Hansen, Mike P. Harris, Martin Irestedt, Oddmund Kleven, Mark L. Mallory, Kjetill S. Jakobsen, Sanne Boessenkool

**Affiliations:** Centre for Ecological and Evolutionary Synthesis (CEES), Department of Biosciences, University of Oslo, Blindernveien 31, 0371 Oslo, Norway; WSL Swiss Federal Research Institute, Zürcherstrasse 111, 8903 Birmensdorf, Switzerland; Norwegian Institute for Nature Research (NINA), Høgskoleringen 9, NO-7034 Trondheim, Norway; Norwegian Polar Institute, Fram Centre, Postbox 6606, Langnes, 9296 Tromsø, Norway; Faroe Marine Research Institute (FAMRI), Nóatún 1, FO-100 Tórshavn, Faroe Islands; Centre for Biodiversity Dynamics (CBD), Norwegian University of Science and Technology (NTNU), Trondheim, Norway; Environment and Climate Change Canada, Newfoundland and Labrador, Canada; Littoral, Environment et Sociétés (LIENSs), UMR 7266 CNRS – La Rochelle Université, 17000 La Rochelle, France; South Iceland Nature Research Centre, Ægisgata 2, 900 Vestmannaeyjar, Iceland; UK Centre for Ecology & Hydrology, Penicuik, Midlothian, EH26 0QB, UK; Department for Bioinformatics and Genetics, Swedish Museum of Natural History, Box 50007, 104 05 Stockholm, Sweden; Department of Biology, Acadia University, Wolfville, Nova Scotia, Canada, B4P 2R6

## Abstract

The factors underlying gene flow and genomic population structure in vagile seabirds are notoriously difficult to understand due to their complex ecology with diverse dispersal barriers and extensive periods at sea. Yet, such understanding is vital for conservation management of seabirds that are globally declining at alarming rates. Here, we elucidate the population structure of the Atlantic puffin (*Fratercula arctica*) by assembling its reference genome and analyzing genome-wide resequencing data of 72 individuals from 12 colonies. We identify four large, genetically distinct clusters, observe isolation-by-distance between colonies within these clusters, and obtain evidence for a secondary contact zone. These observations disagree with the current taxonomy, and show that a complex set of contemporary biotic factors impede gene flow over different spatial scales. Our results highlight the power of whole genome data to reveal unexpected population structure in vagile marine seabirds and its value for seabird taxonomy, evolution and conservation.

## Main

Seabirds are important ecosystem indicators and drivers^e.g. 1–3^, and have long had an integral place in human culture and economy^4–6^. Nevertheless, global seabird numbers have deteriorated by an alarming 70% since halfway of the 20th century^e.g. 7,8^. These declines pose a serious threat to marine ecosystems, human society, and culture^e.g. 7,9,10^, highlighting the importance of seabird conservation management. Within such management the identification of distinct population units is a fundamental first step towards optimized conservation^11–13^. Defining such units is, however, difficult for many seabirds because their complex ecology promotes dispersal barriers due to e.g. oceanography, foraging behavior, at-sea range, and phenology^14^. Detailed genomic data including thousands of loci provide new possibilities to assess levels of connectivity and gene flow between distinct breeding populations and, thus, help identify relevant conservation units for seabirds^14,15^. Indeed, a few recent publications using reduced genomic representation approaches (e.g. RAD-seq) report fine-scale structure over various spatial scales^16–20^. These studies highlight the great potential of genomic data to disentangle barriers to gene flow that would otherwise remain undetected, but remain nonetheless limited due to incomplete sampling of the genome^21^.

The Atlantic puffin (*Fratercula arctica*, Linnaeus, 1789, hereafter “puffin”) is an iconic seabird species, prevalent in popular culture^e.g. 22^, important for tourism^e.g. 23,24^ and inherently valuable for the marine ecosystem^e.g. 1^. Puffins were historically widely harvested for their meat and down^6,25,26^ and exploitation remains an important cultural tradition in Iceland and the Faroe Islands^6,23^. Its breeding range stretches from the Arctic coast and islands of European Russia, Norway, Greenland, and Canada, southward to France and the USA^27^ (Figure 1a). Puffins have been designated as “vulnerable” to extinction globally and listed as “endangered” in Europe^28^. Notably, the once world’s largest puffin colony (Røst, Norway) has experienced complete fledging failure during nine of the last 13 seasons and has lost nearly 80% of its breeding pairs during the last 40 years^28–30^. Similarly, Icelandic and Faroese puffins have experienced low productivity and negative population growth since 2003^31^. Puffins have been broadly classified into three taxonomic groups along a latitudinal gradient based on size, with the *smallest* puffins found around France, Britain, Ireland and southern Norway (*F. a. grabae*), *intermediate* sized puffins around Norway, Iceland and Canada (*F. a. arctica*) and the *largest* puffins found in the High Arctic, e.g. Spitsbergen, Greenland and northeastern Canada^32^ (*F. a. naumanni*)^33^ (Figure 1a). Nevertheless, this broad classification into three subspecies has been controversial^27,34,35^ and the population structure of puffins remains unresolved at all spatial scales^34^. This knowledge gap obstructs efforts towards an assessment of dispersal barriers, limits our understanding of cause-and-effect dynamics between population trends, ecology and the marine ecosystem, and hinders the development of adapted large-scale conservation actions.

**Figure 1:**
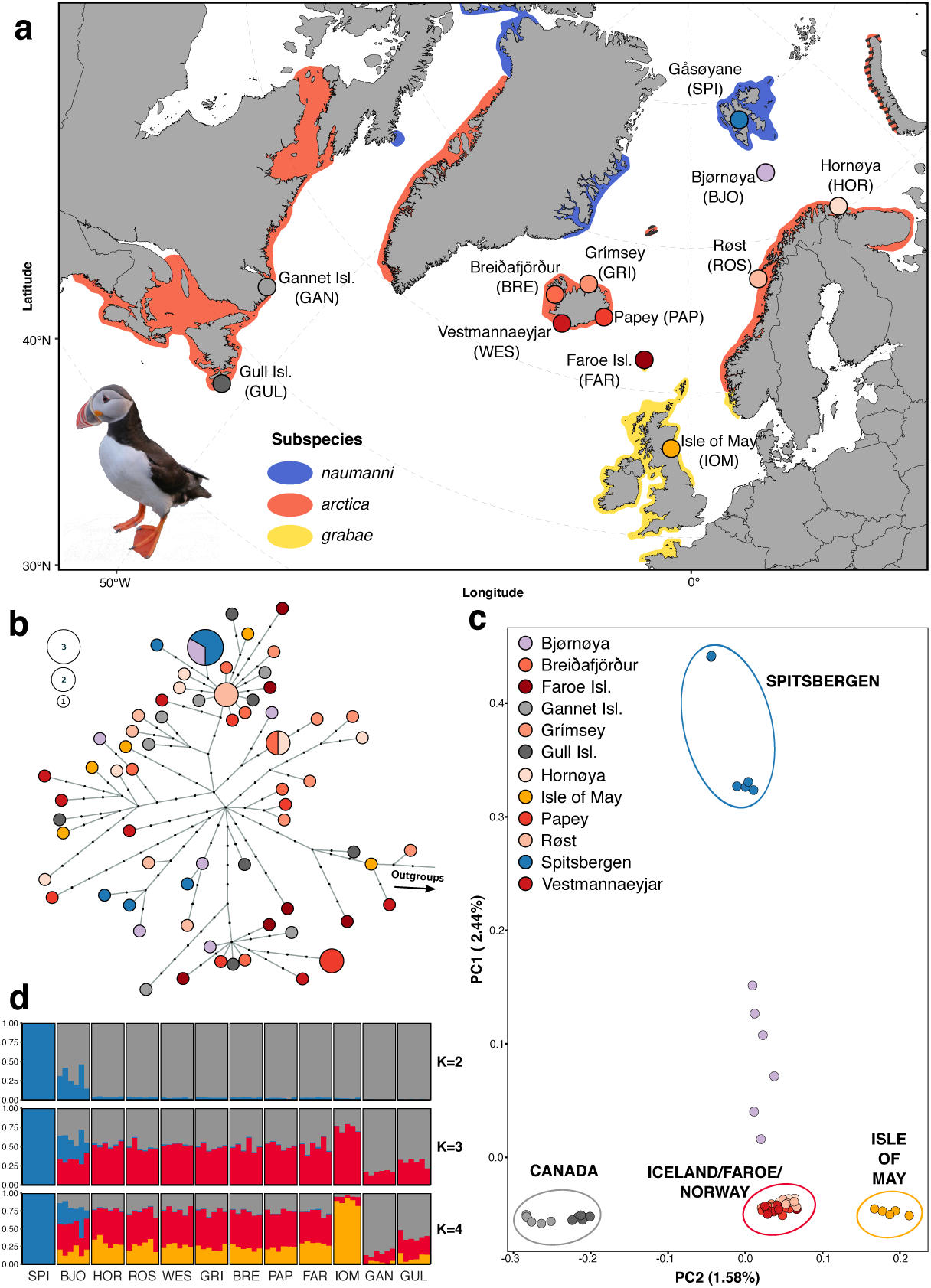
Sampling distribution and genomic structure of 71 Atlantic puffin individuals across 12 colonies throughout the breeding range. a) Map presenting the location of the 12 sampling sites. Colour shading indicates the breeding range of the species as a whole, as well as the recognized subspecies. b) Mitochondrial haplotype network constructed using the program Fitchi and a Maximum Likelihood tree generated in IQTree. It contains 66 unique haplotypes identified by 192 mitogenome-wide SNPs. Sizes of circles are proportional to haplotype abundance. Color legend is provided in c). Black dots represent inferred haplotypes that were not found in the present sampling. c) Principal Component Analysis (PCA) using genotype likelihoods at 1,093,765 polymorphic nuclear sites calculated in ANGSD to project the 71 individuals onto PC axes 1 and 2. Each circle represents a sample and colours indicate the different colonies. Percentage indicates proportion of genomic variation explained by each axis. The colour coding of the colonies is consistently used throughout the manuscript. d) CLUMPAK-averaged admixture plots of the best K’s using the same genotype likelihood panel as in c). Each column represents a sample and colonies are separated by solid white lines. Optimal K’s were determined by the method of Evanno et al. (2005) (see Figure S5a) and colors indicate the ancestry fraction to the different clusters.

Here, we present the first whole genome analysis of structure, gene flow and taxonomy of a pelagic, North Atlantic seabird. We generated a *de novo* draft assembly for the puffin and resequenced 72 individuals across 12 colonies representing the majority of the species’ breeding range (Figure 1a). Our work provides compelling evidence that a complex interplay of ecological factors contributes to the range-wide genomic population structure of this vagile seabird.

## Results

### Genome assembly and population sequencing

Based on synteny with the razorbill (*Alca torda*), a total of 13,328 puffin scaffolds were placed into 26 pseudo-chromosomes, leaving 17.06 Mbp (1.4%) unplaced and yielding an assembly of 1.294 Gbp (Table S1, S2). This assembly contains 4,522 of the 4,915 genes (92.0%) of complete protein coding sequences from the avian set of the OrthoDB v9 database (Table S1). We also assembled the puffin mitogenome (length of 17,084 bp) with a similar arrangement of genomic elements as other members within the bird family Alcidae^36,37^ (Figure S1, Table S3). For the 72 resequenced specimens, we analyzed a total of 5.77 billion paired reads, obtaining an average fold-coverage of 7X (range 3.0 - 10) for the nuclear genome and 591X (5.3 - 1800) for the mitochondrial genome per specimen (Figure 1a, Table S4). One individual (IOM001) was removed from the dataset due to a substantially lower number of mapped reads (endogeny) relative to all other samples (Table S4) resulting in a large proportion of missing sites (Figure S2). Additional filtering produced a final genotype likelihood dataset of 1,093,765 polymorphic nuclear sites and 192 mitochondrial single-nucleotide-polymorphisms (SNPs) in 71 birds (36 males and 35 females).

### Genomic population structure

Genomic variation across 71 puffin mitogenomes defines 66 polymorphic haplotypes that indicate a recent global population expansion and show no significant population structure (Figure 1b, Figure S3, Table S5, S6). In contrast, we inferred four main population clusters using Principal Component Analysis (PCA) of the nuclear whole-genome dataset (Figure 1c). Puffins from Spitsbergen are most distinct, while puffins from Bjørnøya are located between Spitsbergen and a larger, central cluster consisting of populations from Norway, Iceland and the Faroe Islands (Figure 1c, Figure S4a). Puffins from Canada form their own distinct cluster, as do those from the Isle of May, southeast Scotland (Figure 1c, Figure S4b). Hierarchical PCA analyses of the cluster comprising the mainland Norwegian, Icelandic and Faroese colonies reveal further fine-scale structure separating Norwegian (Hornøya and Røst) and Faroese/Icelandic colonies (Figure S4c). Model-based clustering (ngsAdmix) agrees with the results from the PCA (Figure 1b). The optimal model fit for the entire dataset is either K = 2 or K = 4 (Figure S5a), as determined by the method of Evanno et al.^38^. At K = 2, ngsAdmix separates Spitsbergen from the other colonies, with Bjørnøya being admixed (following separation along PCA 1), whereas at K = 4, ngsAdmix reflects the structure of three additional distinct clusters representing Spitsbergen, Canada, the Isle of May, and a central group with more shared ancestry. The shared ancestry of the central group remains present in hierarchical admixture analyses excluding Spitsbergen and Bjørnøya individuals (Figure S5b, S6). We find no fixed alleles and pairwise F_ST_ values between colonies and genomic clusters are low (<0.01) (Table S6), apart from any comparisons involving the Spitsbergen population, which show substantially higher F_ST_ values (0.03-0.08).

Phylogenetic reconstructions using individual-based Neighbor-Joining (NJ) and maximum likelihood (ML) methods (Figure 2a, Figure S7), as well as population-based analyses in Treemix (Figure 2b), support the distinctiveness of the Spitsbergen, Canada and the Isle of May puffins with each group forming monophyletic clades with 100% bootstrap support. In contrast, Bjørnøya forms a paraphyletic clade between Spitsbergen and northern Norway (Figure 2a). The population clusters identified by the PCA and ngsAdmix at smaller spatial scales are also identified in the topologies of the NJ and ML trees, sorting individuals predominantly according to geographical location, although with low bootstrap support (> 80) due to large inter-individual variability (Figure 2a, S6). Allowing a single migration edge in the Treemix phylogeny identifies recent gene flow from Spitsbergen to Bjørnøya (likelihood = 792.106; Figure S8, S9a). Adding additional migration edges to the population-based ML tree does not improve the model fit and such edges are therefore not further interpreted (Figure S8-S10).

**Figure 2:**
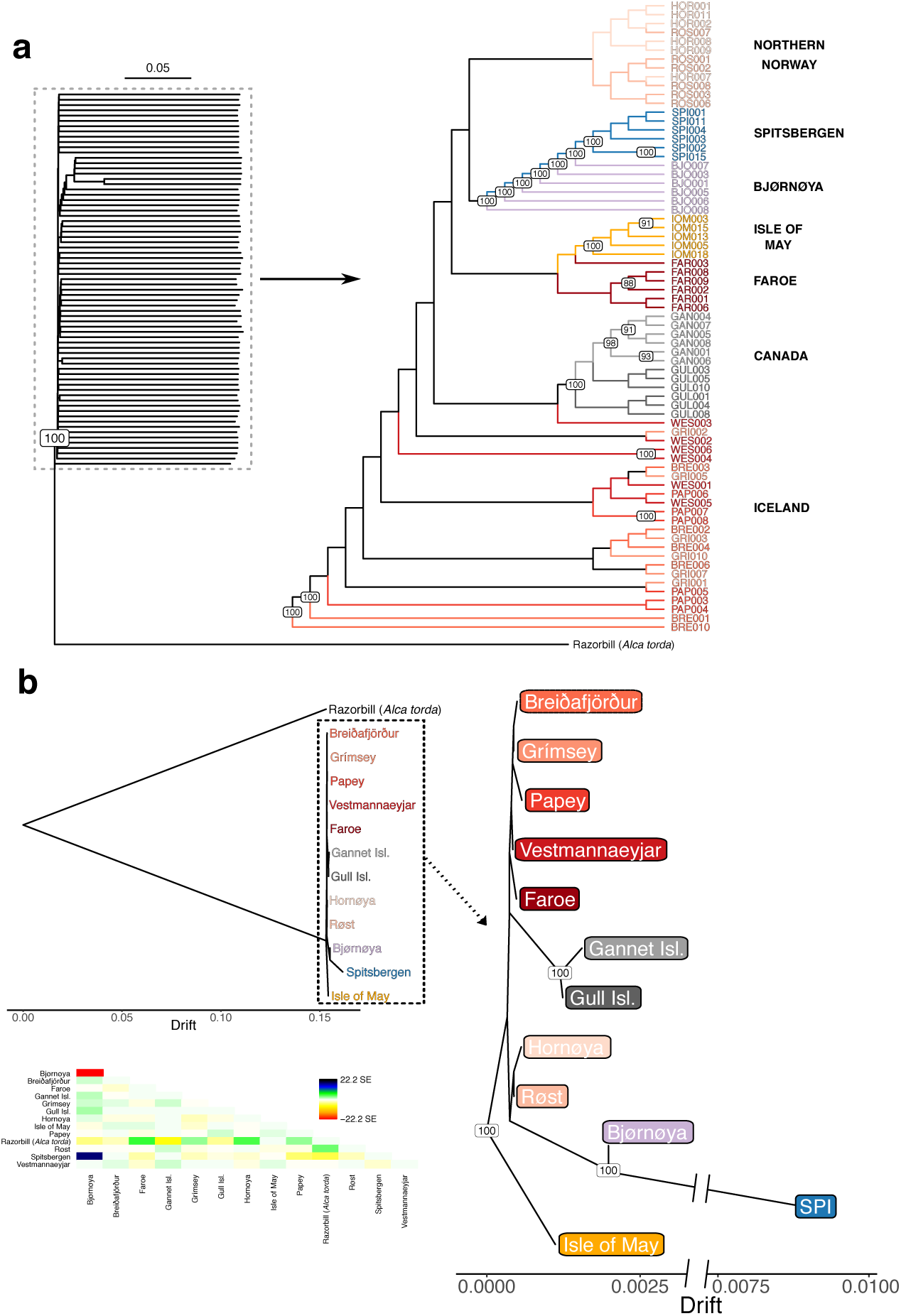
Phylogenetic reconstruction of individual and colony relationships from 71 Atlantic puffin individuals sampled across 12 colonies throughout the species’ breeding range. a) An individual-based neighbor-joining tree constructed using pairwise p-distances calculated from genotype likelihoods at 1,093,765 polymorphic nuclear sites. Branch lengths and the outgroup were removed for the zoomed-in section to improve visualization. b) A population-based maximum likelihood Treemix analysis using allele frequencies at the same 1,093,765 polymorphic nuclear sites as in a). Both trees are rooted using the razorbill as an outgroup. The tree in b) is visualized with and without the outgroup. Branch lengths are equivalent to a genetic drift parameter. The heatmap indicates the residual fit of the tree displaying the standard error of the covariance between populations. In a) and b), the colour coding of the colonies is consistent with those in Figure 1 and node labels show bootstrap support >80.

### Genetic diversity, heterozygosity and inbreeding

Tajima’s D does not significantly deviate from neutral expectation per colony (Table S5). Nucleotide diversity (*π*) of puffins is significantly different between colonies, with the Spitsbergen population having significantly lower nucleotide diversity than the global median (Wilcoxon Rank Sum test, U = 4824, n_SPI_ = 25, n_Global_ = 300, p < 0.05, Table S5). Colonies also differ significantly in levels of heterozygosity (Kruskal-Wallis Test, p < 0.001; Figure 3a) and inbreeding (Kruskal-Wallis Test, p < 0.001, Figure 3b), whereby individual inbreeding (F_RoH_) was approximated based on Runs of Homozygosity (RoH)^39^. Again, the Spitsbergen colony has significantly lower levels of heterozygosity (0.00220 - 0.00223) and significantly higher levels of F_RoH_ values (0.161 - 0.172), compared to the Faroese and Icelandic colonies (Dunn test with Holm correction, p < 0.05). The Faroese and Icelandic colonies contain the highest levels of heterozygosity and lowest F_RoH_ values (Figure 3a, 3b, S11) overall. The remaining colonies display intermediate levels (Figure 3a, 3b), although heterozygosity is significantly lower (Figure 3a, S11) and inbreeding is significantly higher (Figure 3b, S11) on Gull Island and Bjørnøya compared to the Icelandic and Faroese colonies (Dunn test with Holm correction, p < 0.05). Moreover, Spitsbergen harbors the most (an average of 718 per individual) and longest RoHs with eight being ≥ 2.3 Mbp long (4.21 ± 3.02 % of respective chromosome), whereas none of the RoHs in the remaining colonies were > 2.15 Mbp long (Figure 3c). The only exception was a 9.65 Mbp long RoH on pseudo-chromosome 7 (18% of chromosome length) in an Isle of May individual (Figure 3c).

**Figure 3:**
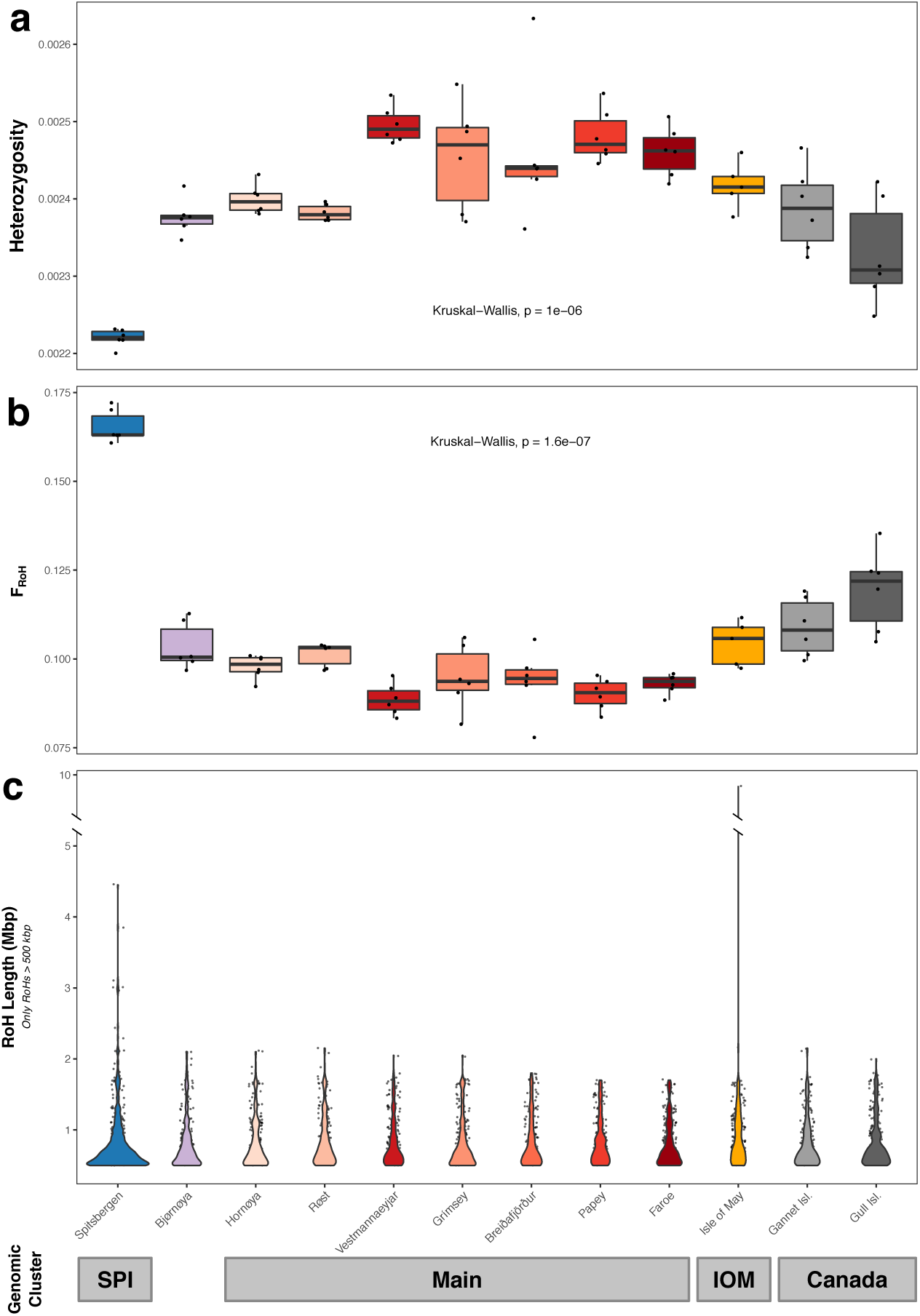
Genome-wide heterozygosity, inbreeding and Runs-of-Homozygosity (RoH) compared between 12 Atlantic puffin colonies across the species’ breeding range. a) Estimates of individual genome-wide heterozygosity based on the per-sample one-dimensional Site Frequency Spectrum calculated in ANGSD. b) Individual inbreeding coefficients, FRoH, defined as the fraction of the individual genomes falling into RoHs of a minimum length of 150 kbp. RoHs were declared as all regions with at least two subsequent 100 kbp windows harboring a heterozygosity below 1.435663 x 10^−3^. c) RoH length distribution across the 12 colonies only including RoHs longer than 500 kbp. A single 9.65 Mbp long RoH on pseudo-chromosome 7 in an Isle of May individual required to introduce a break in the y-axis. In a) and b), black dots indicate individual sample estimates and black lines the median per colony, while in c), black dots represent single RoHs. Statistical significance of differences in heterozygosity and FRoH between populations was assessed with a global Kruskal-Wallis test. The results of *post hoc* Dunn tests with Holm corrections are presented in Figure S11. Different colonies in all three plots are indicated using the same color code as in Figure 1.

### Patterns of gene flow and Isolation-By-Distance (IBD)

We investigated patterns of gene flow and IBD between the colonies using two-dimensional estimated effective migration surface (EEMS) analyses. Levels of gene flow between the Icelandic and Faroese colonies and within the Canadian group is high (3-10x higher than the global average), while intermediate between the Norwegian mainland colonies (around the global average). In contrast, the Spitsbergen colony is split from the remaining colonies by migration rates up to 100x lower than the global average (Figure 4a, S12), while additional regions of low gene flow (2-3x lower than the global average) separate the Isle of May, Canadian, and Bjørnøya colonies from the rest (Figure 4a, Figure S12). Geographic distance between all puffin colonies is a poor predictor of pairwise genetic distance, driven by the high Slatkin’s linearized F_ST_ values of the Spitsbergen colony (Figure S13). Nevertheless, geographic distance among a subset of puffin colonies is significantly correlated with genetic distance using Mantel tests (Figure 4b, S13). Specifically, by progressively removing the more distant colonies (Spitsbergen, Bjørnøya, Isle of May, Canada), the fit of a linear IBD model is significantly improved and the proportion of genetic dissimilarity explained by geographic distance is more than doubled (No SPI = 38.16 %, SPI/BJO/IOM removed = 79.77 %) (Figure 4b, S13). The isolated colonies are characterized by high Slatkin’s linearized F_ST_ values at relatively small geographic distances (Figure S13).

**Figure 4:**
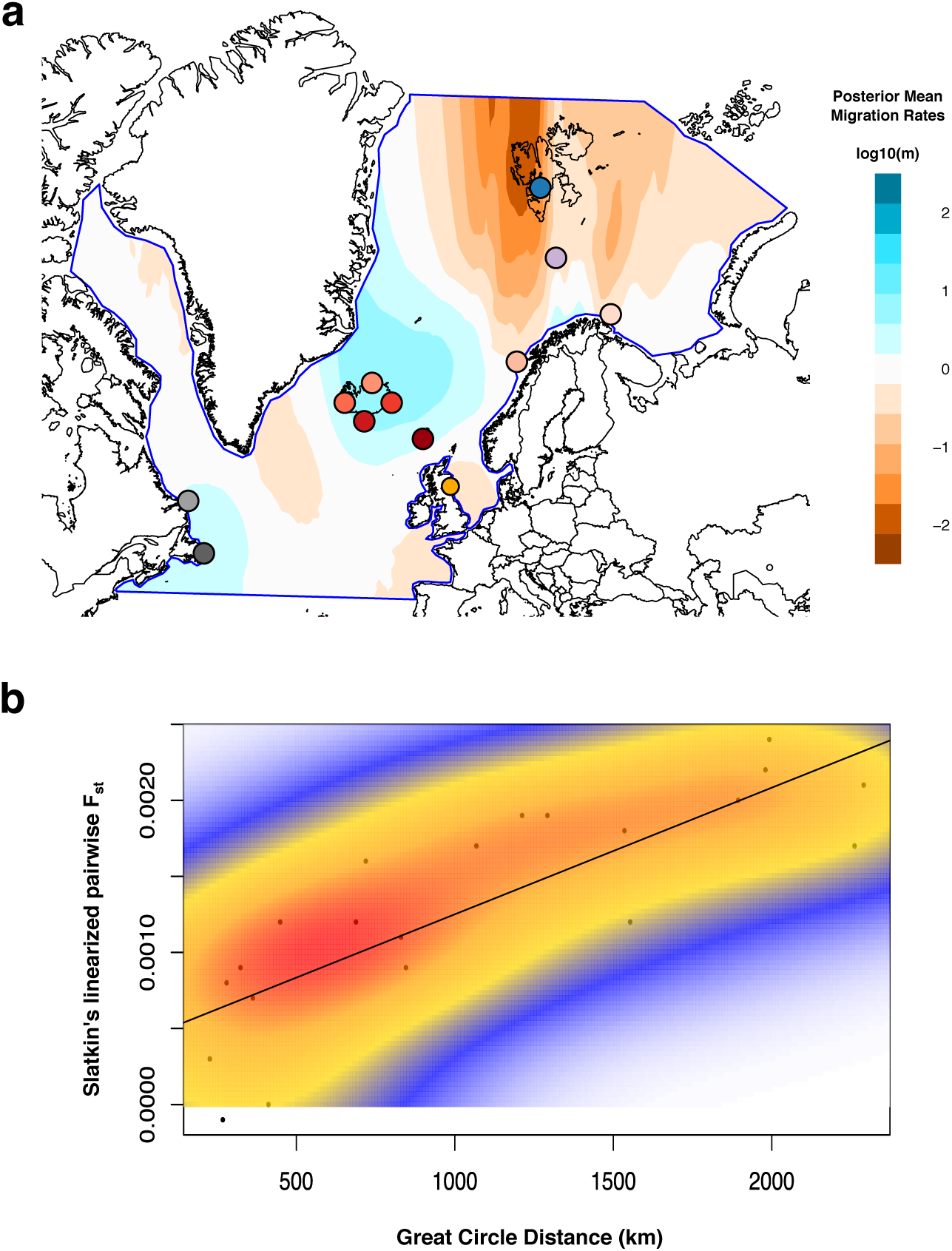
Estimates of continuous long-distance gene flow and isolation by distance (IBD) across the breeding range of the Atlantic puffin estimated from 71 individuals across 12 colonies. a) Effective migration surfaces inferred by the program EEMS using the average distance between pairs of individuals calculated in ANGSD by sampling the consensus base for each individual at 1,093,765 polymorphic nuclear sites. Darker reds indicate reduced migration across those areas, while darker blues highlight higher migration rates than the global mean. Different colonies are indicated using colors consistent with those in Figure 1. b) Correlation between genetic (Slatkin’s linearized FST) and geographic (great-circle) distance presented after removing the Spitsbergen, Bjørnøya, Isle of May and Canadian individuals. The diagonal line visualizes the result of the Multiple Regression on Distance Matrices (MRM) analysis (slope and y-intercept). The Mantel test between genetic and geographic distance (r = 0.826, p < 0.01) was significant and 68.18% of variance in Slatkin’s linearized FST was explained by geographic distance (regression coefficient of linear IBD model = 0.83×10^−6^, p < 0.01). A two-dimensional kernel density estimation (kde2d) highlights dense groups of data points, thus substructure in the genomic landscape pattern. Analyses were conducted and results visualized in R using the *ecodist, marmap* and *MASS* packages.

### Admixture on Bjørnøya

We specifically tested for patterns of admixture in Bjørnøya. Significantly negative *f*3 statistics (Z score < -3) are found for all unique combinations of the phylogeny (Spitsbergen, *X*; Bjørnøya) (Table S7), indicating an admixed colony on Bjørnøya caused by gene flow between Spitsbergen and the remaining colonies. Similarly, significantly positive D-statistics (Z score > 3) caused by an excess of ABBA sites reveal excessive allele sharing between Spitsbergen and Bjørnøya (Figure S14a). The close association and gene flow from Spitsbergen to Bjørnøya is further confirmed by D-statistics not being significantly different from 0 for the (((Bjørnøya, Spitsbergen), H3), Razorbill) topology (Figure S14b).

## Discussion

Barriers to gene flow leading to population structure are notoriously difficult to identify and remain largely unknown for most seabirds^14,40^. Using whole genome analyses, we here provide novel insights into the genetic structure of the Atlantic puffin. First, we identify four main puffin population clusters consisting of (1) Spitsbergen (High Arctic), (2) Canada, (3) Isle of May, and (4) multiple colonies in Iceland, the Faroe Islands and Norway. Second, we find that *within* such clusters, genetic differentiation is driven by isolation-by-distance (IBD). Finally, we find evidence for secondary contact between two clusters. These observations show that a complex set of drivers impacts gene flow over different spatial scales (100s -1000s of km) between these clusters and the colonies within. In particular, the interplay between overwintering grounds, philopatry, natal dispersal, geographic distance, and potentially ocean regimes appears to explain the genomic differentiation between puffin colonies.

Mature puffins rarely, if ever, change their colonies, resulting in very high colony fidelity once they start breeding^27^. Immatures, however, have been observed to visit other nearby colonies during the summer and may breed in non-natal colonies^27,41^. Nevertheless, data on natal philopatry remain scarce, but existing evidence shows rates vary greatly (38-92%) between colonies^27,41^. If either breeding or natal philopatry alone drive the puffin population structure, each colony should constitute its own distinct genomic entity and significant relationships between geographic and genomic distance across the puffin’s entire breeding range would be observed. Yet, philopatry alone cannot explain the presence of the four large-scale population clusters we observe here. Additional factors must therefore promote the distinctiveness of the four clusters. For instance, the Isle of May birds have a largely separate overwintering distribution mainly in the North Sea (Figure S15)^27,35,42^. Such potential geographical separation during the winter season might limit the likelihood of immatures intermixing between the Isle of May and other colonies. Similarly, distinct overwintering distributions have been found to lead to increased genetic diversification in other philopatric seabird species^14,40,43^, such as the thick-billed murre (*Uria lomvia*)^20^ and black-browed albatross (*Thalassarche melanophris*)^44^. The presence of a Canadian cluster can also be largely explained by their winter distribution around Newfoundland^42,45^. There is, however, some fragmentary overlap in the overwintering distribution of the Canadian and Icelandic colonies off southwestern Greenland^42,45^, suggesting that barriers to migration of immatures and gene flow in the western Atlantic may be further enforced by the large geographic distance. In contrast, the winter distribution from the colonies in Iceland, Norway and the Faroe Islands overlaps off the coast of southern Greenland (Figure S15)^42^. This shared overwintering area, combined with the tendency to return to the natal colony and immature visits of nearby (up to 100s km) colonies during the summer, appears to drive a pattern of IBD among colonies (Figure 3b). Indeed, IBD has previously been recognized as an important driver of genomic structure in seabirds, for instance in the little auk (*Alle alle*)^46^ and band-rumped storm-petrel (*Oceanodroma castro*)^47^. While these illustrated mechanisms provide reasonable explanations for the observed dispersal barriers and population structure based on our current knowledge, validation requires additional evidence, specifically on the winter distribution of immature puffins and natal emigration rates across colonies covering the entirety of the puffin’s breeding range.

High Arctic puffins from Spitsbergen are genetically the most divergent group within our dataset. They are also characterized by significantly lower levels of genetic diversity, greater inbreeding coefficients, and longer and more abundant RoHs compared to other colonies. These observations may either result from a historical bottleneck followed by isolation (e.g. founder effect), local adaptation to their extreme environment or generally lower effective population sizes. Population abundance estimates of less than 10,000 breeding pairs on Spitsbergen compared to 500,000 in the West Atlantic, two million on Iceland and more than two million in the boreal East Atlantic potentially indicate a lower effective population size^27^. The High Arctic puffins exclusively inhabit harsh, cold-current environments, also in the winter, during which they likely stay in an area bounded by the East Greenland ice edge, a latitudinal border at 70° N and the front between the Barents and Greenland Sea (Figure S15). They are also substantially larger than birds from lower latitudes^27^, following Bergmann’s^48^ or James’s^49^ rule, as has been observed in other seabirds^e.g. 50,51^. This matches the clinal size variation of puffins that closely tracks sea temperatures in their breeding areas^52^. Despite these distinctions, we find that the relatively small population of puffins on Bjørnøya (< 1,000 pairs^27^), midway between Spitsbergen and mainland Norway, represents an area of secondary contact between the puffins from the High Arctic and other puffin colonies. Based on D- and the *f*3-statistics, the most likely southern sources are Iceland, the Faroe Islands, Norway, or a combination thereof. Thus, the barriers to gene flow that keep the Spitsbergen colonies distinct do not prevent formation of a hybrid colony where individuals from the High Arctic and the cluster composed of mainland Norwegian, Icelandic and Faroese colonies meet.

The distinct population structure in the nuclear data is not observed in the mitochondrial genomes, which reveal an abundance of rare alleles and lack of significant population differentiation. The mitogenomic variation suggests that puffins experienced a recent population expansion, possibly out of a refugium after the Last Glacial Maximum. Indeed, it has been shown that mitogenomic variation in seabirds is dominated by historical factors rather than representing contemporary gene flow^40^, and a lack of mitogenomic population structure has been observed in many marine birds with high philopatry^e.g. 46,53,54^. In contrast to the mitogenomes, the structure in the nuclear data therefore likely originated after the last glacial period and reflects the influence of relatively recent barriers to gene flow in a context of historical demography^14,40^. Such results are relevant for understanding the “seabird paradox”, which contrasts the life-history traits of high philopatry and restricted dispersal in otherwise highly mobile species^55^.

Our results have major implications for the conservation management of the Atlantic puffin. The genetic structure we identify in puffins disagrees with the suggestion of three subspecies (*F. a. naumanni, F. a. arctica, F. a. grabae*)^33^. Although the genetically distinct Spitsbergen cluster coincides with the classification of morphologically large puffins in the High Arctic (*F. a. naumanni*)^27^, we observe gene flow from Spitsbergen into Bjørnøya, which has been considered *F. a. arctica*^*27*^. Furthermore, the geographic divide between *F. a. grabae* and *F. a. arctica* lies farther south than previously thought, with the Faroese puffins being genetically closer to *F. a. arctica* than to *F. a. grabae*. Nonetheless, *F. a. grabae* is currently represented by a single colony (Isle of May) in our study and the geographical extent of this genomic cluster needs to be refined by additional sampling, particularly in the western UK, Ireland, and France. Finally, puffins from the Western Atlantic region (e.g. colonies in Canada) form their own distinct genetic cluster that is not recognized within the current classification. Our results do not only warrant a revision of Salomonsen’s taxonomic classification of three subspecies^33^, but also highlight the need to acknowledge the four identified clusters as distinct units within the conservation management of puffins^11–13^. Although puffin colonies *within* clusters are not genetically distinct entities, ecological independence illustrated by contrasting population dynamics across relatively small spatial scales (e.g. western Norway^30^) suggests that higher resolution local management units based on ecological differences should be considered. Nonetheless, the observation that the genetically distinct clusters are at the outer edges of the puffin’s distribution, indicates that the management and conservation of these distinct clusters must be a priority. Finally, our sampling does not cover several outskirts of the puffin’s distribution, such as the US, northern Canada, Greenland, Ireland, western UK, France or Russia, and we may therefore still underestimate the true biological and genetic complexity of this species.

In conclusion, our study shows that a complex interplay of barriers to gene flow drives a previously unrecognized population diversification in the iconic Atlantic puffin. So far, much of seabird population genetics research has been based on mitochondrial and microsatellite data^14,40^, which have limited power to characterize contemporary factors that determine population structure and gene flow^19,56^. High-resolution nuclear data are therefore essential to help define evolutionary significant population units, disentangle convoluted ecological relationships, and are particularly important for seabird conservation, which aims to preserve genetic diversity considering profound global population declines^e.g. 7,8^ and the threat of global warming, which negatively impacts ecosystems worldwide^57^.

## Methods

### Draft reference genome assembly

A *de novo* Atlantic puffin draft genome was generated from the blood of a female Atlantic puffin. Read data was sequenced on three Illumina HiSeqX lanes using the 10x Genomics Chromium technology and assembled with the Supernova assembler (v2.1.1)^58^ after subsampling to 0.8 billion and 1 billion reads to maximize performance and remain within the computational capacity of the assembler. We refined the two assemblies through several steps, including merging of ‘haplotigs’, removal of contaminant sequences, misassembly correction, re-scaffolding using mapping coverage and linkage information, and gap filling (Table S1a). The most complete and continuous 800M and 1000M assemblies together with the 3rd best assembly overall were selected for a second round of refinement (Table S1b) resulting in a total of 72 draft assemblies. Of these, we kept the four most complete and continuous assemblies for additional gap filling and polishing, after which the most complete draft genome was selected for downstream analyses (Table S1c). The puffin mitogenome was confidently identified by blasting (blastn) all scaffolds shorter than 25 kb against a custom-built database of 135 published mitogenomes of the order ‘Charadriiformes’ and annotated with the MITOS web server^59^ (Figure S1). The remaining nuclear scaffolds were ordered and concatenated into “pseudo-chromosomes’’ by mapping them to the razorbill genome (*Alca torda* - NCBI: bAlcTor1 primary, GCA_008658365.1) and applying 200 N’s as padding between each scaffold. We combined unmapped scaffolds into an “unplaced” pseudo-chromosome. We assessed order and placement of scaffolds by investigating synteny in coverage and length between the puffin and razorbill chromosomes (Table S2). Details on the draft reference genome assembly and refinement can be found in the Supplementary File.

### DNA extraction and sequencing

Samples from a total of 72 puffins collected across 12 breeding colonies (Figure 1a) were made available for the present study by SEAPOP (http://www.seapop.no/en), SEATRACK (http://www.seapop.no/en/seatrack) and ARCTOX (http://www.arctox.cnrs.fr/en/home - Canadian colonies). These samples had been collected between 2012-2018 and consisted of blood preserved in EtOH or lysis buffer, or feathers (Table S4). We extracted DNA using the DNeasy Blood & Tissue kit (Qiagen) following the manufacturer’s protocol for animal blood or the nail/hair/feathers protocol applying several modifications for improved lysis and DNA yield. Individuals that had no sexing data associated with them were sexed using PCR amplification of specific allosome loci and visualization via gel electrophoresis. Genomic libraries were built by the Norwegian Sequencing Centre and sequenced on an Illumina HiSeq4000. We processed sequencing reads in PALEOMIX v1.2.14^60^ and split the resulting bam files into nuclear and mitochondrial bam files. Additional details on the DNA extraction, sexing, sequencing and mapping are listed in the Supplementary File.

### Mitogenome analyses

Genotypes across the mitochondrial genome were jointly called with GATK v4.1.4^61^ by using the *HaplotypeCaller, CombineGVCFs* and *GenotypeGVCFs* tool. We filtered genotypes according to GATKs Best Practices^62^ and set genotypes with a read depth less than 3 or a quality less than 15 as missing. Indels and non-biallelic SNPs were removed and only SNPs present in all individuals were kept for subsequent analyses. The SNP dataset was annotated with snpEff^63^ utilizing the annotation of the newly assembled mitogenome of the Atlantic puffin and converted into a mitogenome sequence alignment. To serve as an outgroup, we appended four other species of the family Alcidae, i.e. the Razorbill (*Alca torda*, NCBI: CM018102.1), the Crested Auklet (*Aethia cristatella*, NCBI: NC_045517.1), the Ancient Murrelet (*Synthliboramphus antiquus*, NCBI: NC_007978.1) and the Japanese Murrelet (*Synthliboramphus wumizusume*, NCBI: NC_029328.1), to the alignment. To construct a maximum-likelihood phylogenetic tree, we split the alignment into seven partitions, i.e. one partition for a concatenated alignment of each of the three codon positions of the protein coding genes, one partition for the concatenated alignment of the rRNA regions, one partition for the concatenated alignment of the tRNAs, one partition for the alignment of the control region, and one partition for the concatenated alignment of the “intergenic” regions. The best-fitting evolutionary model for each partition was found by *ModelFinder*^*64*^ and the tree was built with IQTree v1.6.12^65^ using 1000 ultrafast bootstrap replicates. We used the resulting tree to draw a haplotype genealogy graph with Fitchi^66^. Using Arlequin v.3.5^67^, we calculated haplotype (h), nucleotide diversity (π) and Tajima’s D^68^ for each colony, for each genomic cluster defined by the nuclear analysis, and globally. Additionally, an Ewens–Watterson test^69^, Chakraborty’s test of population amalgamation^70^ and Fu’s F_s_ test^71^ were conducted for each of those groups. To further identify population differentiation, the proportion of sequence variation (Φ_ST_) was estimated for all pairs of populations and genomic clusters. Hierarchical AMOVA tests subsequently determined the significance of *a priori* subdivisions into colonies and genomic clusters. Calculation of Φ_ST_ and AMOVA tests were also conducted in Arlequin. Additional details on the mitochondrial analyses are given in the Supplementary File.

### Nuclear genome clustering and phylogenetic analyses

The majority of population genomic analyses were based on nuclear genotype likelihoods as implemented in ANGSD v.0.931^72^. After assessing the quality of the mapped sequencing data in an ANGSD pre-run, we removed an individual from the Isle of May from the dataset. Genotype likelihoods for nuclear SNPs covered in all individuals were calculated and filtered in ANGSD. Accounting for linkage disequilibrium, we further pruned the dataset by only selecting the most central site within blocks of linked sites (r^2^>0.2) as in Orlando and Librado (2019)^73^. Subsequently, all variants located on the Z-pseudo-chromosome and “unplaced scaffolds” were excluded from the analyses yielding a final genotype likelihood panel consisting of 1,093,765 sites. We investigated genomic population structure with a Principal Component Analysis (PCA) of the genotype likelihood panel using PCAngsd v0.982^74^ and estimated individual ancestry proportions using a maximum likelihood (ML) approach implemented in ngsAdmix v32^75^, with the number of ancestral populations (K) set from 1 to 10. Additional “hierarchical” PCA and admixture analyses were conducted for genomic sub-cluster(s) using identical methods.

After adding the razorbill genome as an outgroup to the genotype likelihood panel by mapping unpublished, raw 10x Genomics sequencing data used for the assembly of the embargoed razorbill genome to the puffin draft assembly, we built a neighbor-joining (NJ) tree based on pairwise genetic distance matrices (p-distance) and a sample-based ML phylogenetic tree in FastMe v2.1.5^76^ and Treemix v1.13^77^, respectively. For both trees, 100 bootstrap replicates were generated. To infer patterns of population splitting and mixing, we produced population-based ML trees including up to ten migration edges. The optimal number of migrations was selected using a quantitative approach by evaluating the distribution of explained variance, the log likelihoods, the covariance with an increase in migration edges, and by applying the method of Evanno^38^ and several different linear threshold models. The topology for m_0_ and m_BEST_ was evaluated by generating 100 bootstrap replicates. Additional details on the cluster and phylogenetic analyses are given in the Supplementary File.

### Genetic diversity, heterozygosity and inbreeding

We calculated a set of neutrality tests and population statistics in ANGSD using colony-based one-dimensional (1D) folded Site-Frequency-Spectra (SFS). For each population, genomic cluster, and globally, Tajima’s D and nucleotide diversity (Π) were computed utilizing the per-site Πestimates. Individual genome-wide heterozygosity was calculated in ANGSD using individual, folded, 1D SFS. We calculated heterozygosity by dividing the number of polymorphic sites by the number of total sites present in the SFS.

The proportion of runs of homozygosity (RoH) within each puffin genome was computed by calculating local estimates of heterozygosity in 100 kbp sliding windows (50 kbp slide) following the approach in Sánchez-Barreiro et al. (2020)^39^. We defined the 10% quantile of the average local heterozygosity across all samples as the cutoff for a “low heterozygosity region”. RoHs were declared as all regions with at least two subsequent windows of low heterozygosity (below cutoff) and their final length was calculated as described in Sánchez-Barreiro et al. (2020)^39^. We calculated an individual inbreeding coefficient based on the RoH, F_RoH_, as in Sánchez-Barreiro et al. (2020)^39^ by computing the fraction of the entire genome falling into RoHs, with the entire genome being the total length of windows scanned. Additional details on these analyses can be found in the Supplementary File.

### Patterns of gene flow and admixture

Assessing potential landscape genetic patterns of IBD within the breeding range of the puffin, the program EEMS^78^ was used to model the association between genetic and geographic data by visualizing the existing population structure and highlighting regions of higher-than-average and lower-than-average historic gene flow. We calculated a pairwise genetic distance matrix in ANGSD by sampling the consensus base (*-doIBS 2 -makeMatrix 1*) at the sites included in the genotype likelihood set (see *Nuclear cluster and phylogenetic analyses*) for each sample. The matrix was fed into 10 independent runs of EEMS, each consisting of one MCMC chain of six million iterations with a two million iteration burn-in, 9999 thinning iterations, and 1000 underlying demes. Supplementing the results of the EEMS analysis, we conducted a traditional IBD analysis by determining geographical and genetic distances between the 12 colonies and assessing the significance of the correlation between the two distance matrices with a Mantel test^79^ and a Multiple Regression on distance Matrix (MRM)^80^ analysis. F_ST_ was used as a proxy for genetic distance and computed for each population pair in ANGSD by applying two-dimensional (2D), folded SFS. We converted pairwise F_ST_ values to Slatkin’s linearized F_ST_^81^. Great circle distances between colony coordinates (latitude/longitude) were calculated using the R package marmap^82^ and used as geographic distances. We performed the Mantel test (10,000 permutations) and MRM analysis with the R package *ecodist*^*83*^. All analyses for IBD were re-run on subsets of colonies by progressively removing the Spitsbergen, Bjørnøya, Isle of May, and Canadian colonies from the geographic and genetic distance matrices.

Additional assessments of gene flow and admixture were conducted by calculating *f*3-statistics and multi-population D-statistics (aka ABBA BABA test)^84^. We calculated *f*3-statistics in Treemix for each unique combination of ((A,B),C)) of the 12 puffin populations. D-statistics were calculated in ANGSD (-doAbbababa2) for each combination of ((A,B),C),Outgroup) using the 12 puffin colonies. The outgroup was generated in ANGSD using the 10xGenomics sequencing data of the razorbill mapped to the puffin reference genome (see *Nuclear cluster and phylogenetic analyses)*. Additional details on the IBD and admixture analyses are given in the Supplementary File.

## Acknowledgements

Financial support was provided by the Nansen Foundation and the Faculty of Mathematical and Natural Sciences, University of Oslo (UiO). We are grateful to the SEAPOP program (www.seapop.no/en, Norwegian Research Council grant number 192141), and the SEATRACK (http://www.seapop.no/en/seatrack) and ARCTOX (https://arctox.cnrs.fr/en/home) projects for collecting and sharing of samples. In particular, we thank Dave Fifield and Greg Robertson for the provision of puffin samples from Gull Island. The authors acknowledge support from the National Genomics Infrastructure in Stockholm funded by the Science for Life Laboratory, the Knut and Alice Wallenberg Foundation and the Swedish Research Council, and the SNIC/Uppsala Multidisciplinary Center for Advanced Computational Science for assistance with massively parallel sequencing (reference genome) and access to the UPPMAX computational infrastructure. Special thanks goes to the Norwegian Sequencing Centre (UiO) for the genomic libraries and resequencing of samples analyzed in this study. The razorbill genome data was made available for this study by Tom Gilbert and the Vertebrate Genome Project. Computation was performed using the resources and assistance from SIGMA2. Albína Pálsdóttir shared scripts for ANGSD, and Emiliano Trucchii advised on the manuscript. Pictures of puffins used in the figures were taken by Annemarie Look.

## Author Contributions

SB and BS conceptualized the project. MI performed the DNA extraction for the reference genome. OK did all other laboratory work. KSJ advised on the sequencing strategy and co-supervised OK. OK refined the reference genome assembly. OK carried out the population genomic analyses with input from SB, BS and DML. OK designed the figures with input from SB, BS and DML. TAN, HS and ESH advised on colony selection and provided ecological context. TAN, HS, OdK, SD, ESH, JF, MPH, JD, KE, MLM and MGF provided samples. OK wrote the paper with SB, BS and DML. All authors read and revised the final version of the manuscript.

## Data availability

Raw read data have been deposited in the European Nucleotide Archive (ENA, www.ebi.ac.uk/ena) under study accession number PRJEB40631 (see Table S4 for individual sample accession numbers). Nuclear and mitochondrial scaffolds (GCA_905066775.1, CAJHIB010000001-CAJHIB010013329), as well as pseudo-chromosomes (GCA_905066775.2, CAJHIB020000001-CAJHIB020000027), have been uploaded to ENA (Project PRJEB40926, Sample SAMEA7482542).

## Code availability

Full code used for the population genomic analyses is available on the first author’s GitHub (https://github.com/OKersten/PuffPopGen).

